# Multiparameter flow cytometric detection and analysis of rare cells *in in vivo* models of cancer metastasis

**DOI:** 10.1101/2023.10.12.562110

**Authors:** Mikaela M Mallin, Kenneth J. Pienta, Sarah R. Amend

## Abstract

Rapid and reliable circulating tumor cell (CTC) and disseminated tumor cell (DTC) detection forms a major underpinning of rigorous *in vivo* metastasis research. While many cancer cells initiate metastasis, very few can complete it. Clinical data evidences that each successive step of metastatic cascade presents increasing barriers to metastatic success, limiting the number of successful metastatic cells to fewer than 1 in 1,500,000,000. As such, it is critical to employ metastasis research approaches that allow scientists to discern which step(s) of the cascade present metastatic barriers to their model systems, and in which steps their model systems might display competency. Here, we present a novel flow-cytometry based method that allows for the simultaneous comparison of multiple steps of the cascade within one model system via the co-identification of CTC and DTCs from single animals. This approach is not only highly reliable and reproducible, but also broadly applicable and highly adaptable to a wide range of scientific inquiries.

## Introduction

The rapid and reliable identification of circulating and disseminated tumor cells from murine tissues is essential to understanding the metastatic potential of cancer cells following experimental manipulation.

The well-described metastatic cascade consists of five distinct steps: invasion by motile cells in the primary tumor site, intravasation into vasculature, survival within the circulatory system as circulating tumor cells (CTCs), extravasation into a distant secondary site as disseminated tumor cells (DTCs) and eventual colonization into a detectable macrometastatic lesion. While successful completion of the entire metastatic cascade appears common at the patient level, complete metastatic-competency is surprisingly rare event at the cellular level. Many barriers to metastasis compound at every step of the cascade, limiting a cell’s metastatic success. Calculations using clinical data (i.e. frequencies of detectable CTCs, DTCs, and outgrown metastatic lesions in patients) indicate that the likelihood of any given CTC to successfully produce a metastatic tumor is less than 1 in nearly 1,500,000,000. A more nuanced understanding of where and when cancer cells that have initiated the metastatic process fail (or much more rarely, succeed) is crucial for the development of metastasis-intervention therapies.

As such, it is critical to employ metastasis research approaches that allow scientists to discern which step(s) of the cascade present metastatic barriers to their model systems, and in which steps their model systems might display competency. A method that allows for the simultaneous comparison of multiple steps of the cascade within one model system is the most optimal approach, such as simultaneous detection of CTCs and DTCs from a single animal. We developed an optimized flow-cytometry-based CTC and DTC detection technique for the identification of metastasizing cells in murine tissues. Our approach utilizes human cancer cell lines transduced to constitutively express both cytosolic Green Fluorescent Protein (GFP) and Luciferase enzyme at high levels. Cancer cells were cultured, injected, and monitored using luciferin-based bioluminescent imaging. At endpoint, metastasis-relevant tissues were collected and processed for same-day flow cytometry analysis, using presence of GFP as basis for identification of CTCs and DTCs.

As proof of concept, we highlight here the successful identification of mouse CTCs and bone marrow DTCs recovered 6 weeks after subcutaneous injection of PC3-GFP-Luc human prostate cancer cells into 10-week old male immunocompromised mice. We also demonstrate the wide applicability of this approach, showing the subsequent successful identification of DTCs resulting from injected high-ploidy cells that were a) cultured to be much larger in size, b) inoculated into the animal using a tail-vein injection technique, and c) recovered from lung tissue, a solid organ tissue. Furthermore, we illustrate the adaptability of this approach, offering an example of the additional fluorophore-based characterizations that can be multiplexed into the experimental design to further define and describe recovered CTCs and DTCs.

## Materials and Methods

Broadly, this process consists of 7 steps: Cell culture, injection of cultured cells, monitoring of injected animals, metastasis-relevant tissue collection, harvested tissue processing for flow cytometry, flow cytometry data collection, and data analysis (Figure 1). The line of cells, injection route, and tissue-type collected are experiment-specific and easily modifiable. Here, we present specific protocols for a) culture of normal and/or of high-ploidy PC3-GFP-Luc prostate cancer cells, b) injection via subcutaneous and/or tail vein route, and c) collection and processing of blood, bone marrow, and/or lung tissues. Due to the live-cell nature of the flow cytometry, tissue collection, processing and data collection must occur on the same day for each cohort of mice. A summary of the necessary materials and equipment is detailed in Table 1. Troubleshooting guidelines are detailed in Table 2.

**Table 2:**
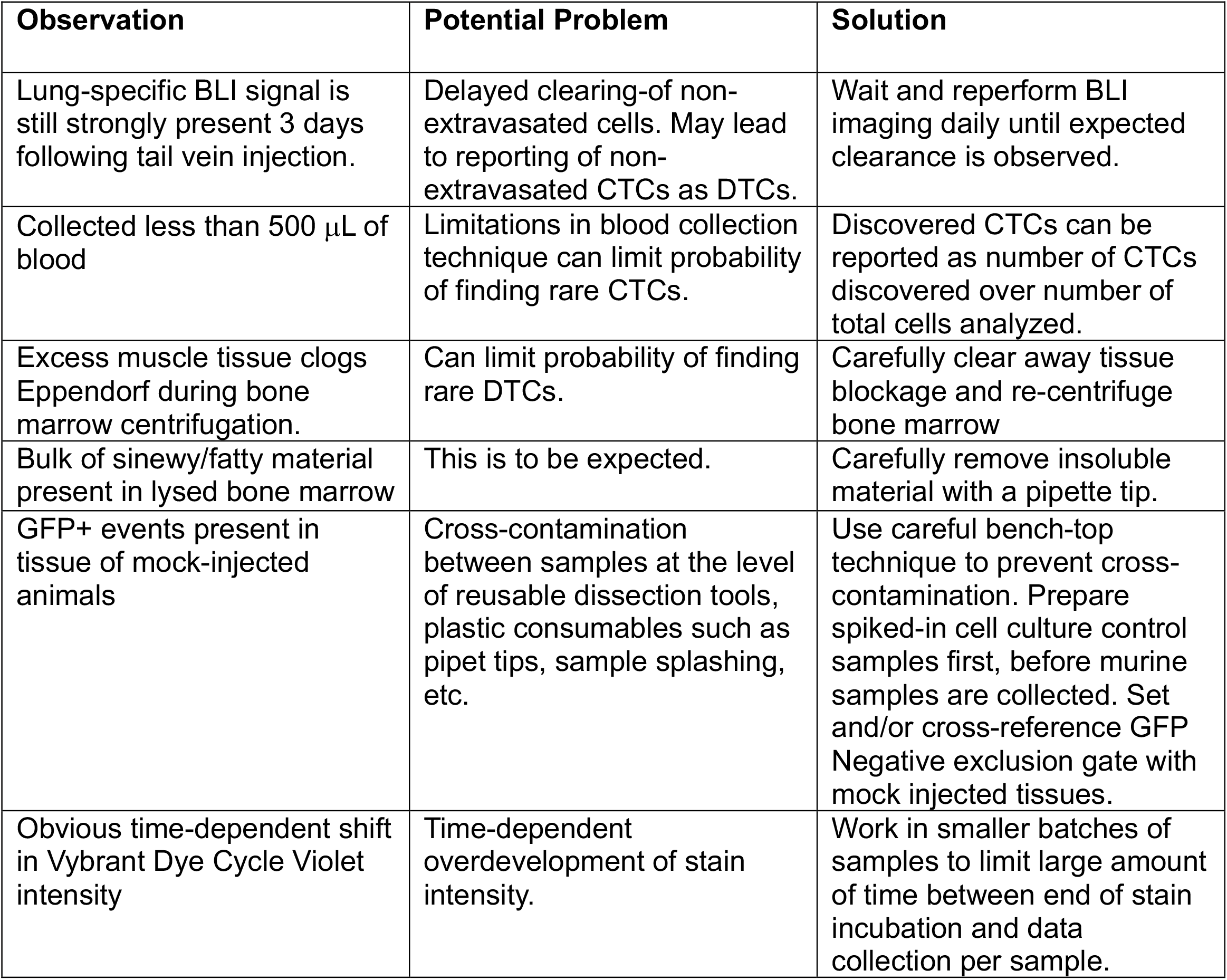
Troubleshooting Guidelines.

| **Observation** | **Potential Problem** | **Solution** |
| --- | --- | --- |
| Lung-specific BLI signal is still strongly present 3 days following tail vein injection. | Delayed clearing-of non-extravasated cells. May lead to reporting of non-extravasated CTCs as DTCs. | Wait and reperform BLI imaging daily until expected clearance is observed. |
| Collected less than 500 μL of blood | Limitations in blood collection technique can limit probability of finding rare CTCs. | Discovered CTCs can be reported as number of CTCs discovered over number of total cells analyzed. |
| Excess muscle tissue clogs Eppendorf during bone marrow centrifugation. | Can limit probability of finding rare DTCs. | Carefully clear away tissue blockage and re-centrifuge bone marrow |
| Bulk of sinewy/fatty material present in lysed bone marrow | This is to be expected. | Carefully remove insoluble material with a pipette tip. |
| GFP+ events present in tissue of mock-injected animals | Cross-contamination between samples at the level of reusable dissection tools, plastic consumables such as pipet tips, sample splashing, etc. | Use careful bench-top technique to prevent cross-contamination. Prepare spiked-in cell culture control samples first, before murine samples are collected. Set and/or cross-reference GFP Negative exclusion gate with mock injected tissues. |
| Obvious time-dependent shift in Vybrant Dye Cycle Violet intensity | Time-dependent overdevelopment of stain intensity. | Work in smaller batches of samples to limit large amount of time between end of stain incubation and data collection per sample. |

**Figure 1:**
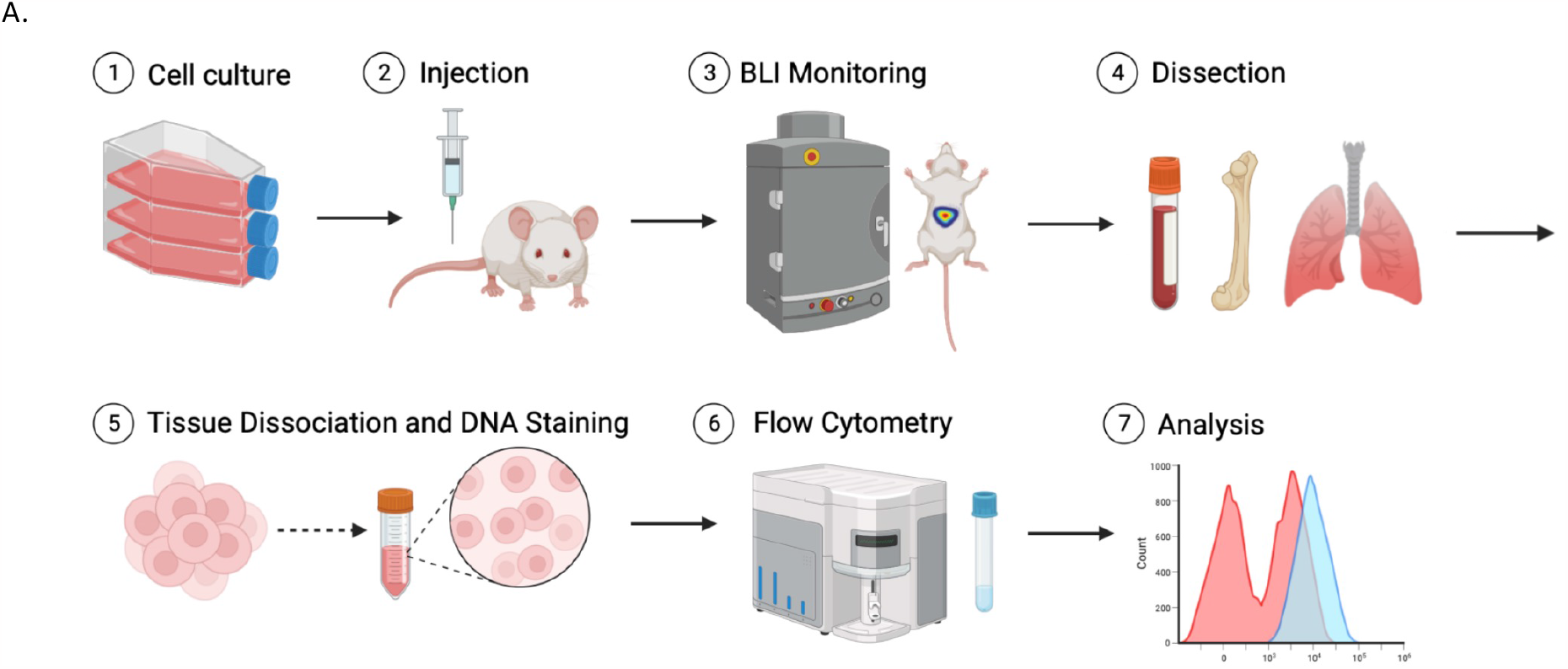
Workflow Schematic A) Graphical abstract outlining the general steps of our featured technique, including cell culture, injection of cultured cells, monitoring of injected animals, metastasis-relevant tissue harvesting, tissue processing, flow cytometry-based data collection, and data analysis.

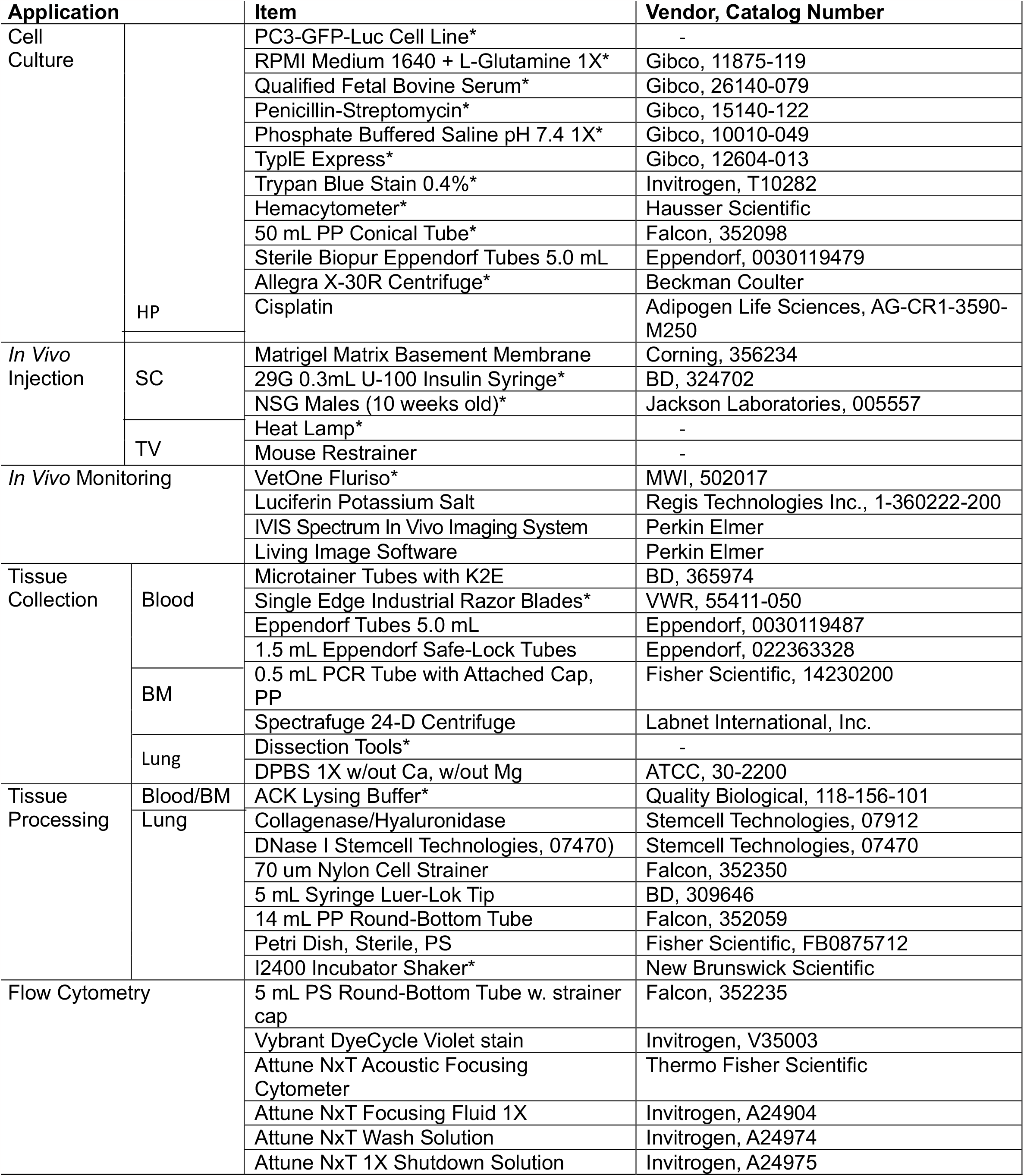
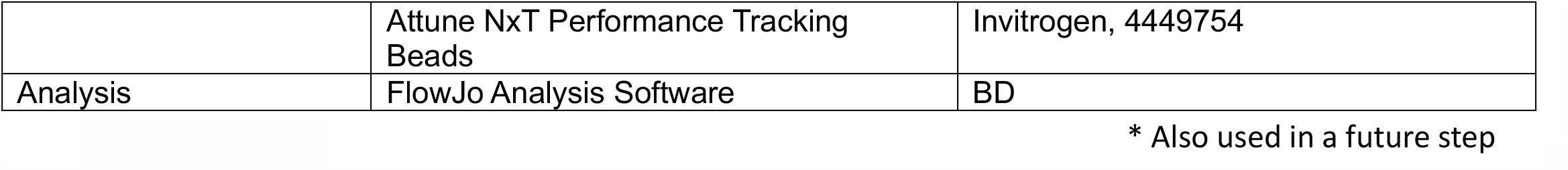

### Cell Culture

#### PC3-GFP-Luc

Experiments were performed with a PC3-GFP-Luc prostate cancer cell line. The cell line was generated from standard PC3 cells (ATCC, CRL-1435) + pLentilox-EV-Luc luciferase expression vector (University of Michigan Vector Core), + pLenti-CMV-GFP-Neo (657-2) GFP expression vector (Addgene, Plasmid 17447). All cells were cultured with RPMI 1640 media, supplemented with 10% Fetal Bovine Serum and 1% Penicillin-Streptomycin antibiotic at 37 degrees Celsius and in 5% CO2. Cells were STR-profile authenticated and tested for *mycoplasma* contamination biannually (Genetica) TryplE Express (following one Phosphate-Buffered Saline wash) was routinely used as the cell dissociation reagent. All centrifugations of cells were performed at 1000 RPM for 5 minutes.

#### High-Ploidy PC3-GFP-Luc

When applicable, high-ploidy PC3-GFP-Luc cells were generated using the following protocol. To accommodate for chemotherapy-induced apoptosis among developing high-ploidy cells, scale tissue culture flask numbers accordingly. Cells were treated with 12 μM of cisplatin for 72 hours. After 72 hours, drug-containing media was removed and and replaced with fresh complete media and cells were cultured for an additional 10 days.

### *In Vivo* Injection

#### Subcutaneous Tumor Injection

100 μL of cells (200,000 cells/mouse) or mock suspension (1:1 v:v Matrigel) were injected subcutaneously into the right flanks of 10-week old male NSG mice.

#### Tail Vein Injection

100 μL of cells (100,000 cells/mouse) or mock suspension were injected into the lateral tail vein of 10-week old male NSG mice

### *In Vivo* Monitoring

Bioluminescent monitoring is used to ensure correct injection technique and appropriate model progression. Subcutaneously injected mice were imaged weekly to track cancer cell growth kinetics. Tail Vein injected mice were be imaged within 30 minutes of injection to confirm correct injection technique via presence of strong lung signal. Tail Vein injected mice should be imaged again immediately prior to tissue collection to confirm attrition of lung-filtered cells that failed to extravasate into lung tissue. Here, a large decline in lung signal should be expected.

### Tissue Collection

#### Blood

Blood was collected at time of sacrifice via tail bleed 6 weeks following subcutaneous injection. Blood was collected directly into K2EDTA treated microtainers (a minimum of 250 μL is necessary for analysis). suffice. Samples were stored on ice until processing.

#### Bone Marrow

Bone marrow was collected as previously described (doi: 10.3791/53936) at 6 weeks post-tumor subcutaneous tumor injection. Samples were stored on ice until processing.

#### Lung

Lung tissue was collected 3 3 days following tail vein injection. Harvest all 5 lobes of the heart. Transfer to PBS+2%FBS to cover the sample. Samples were stored on ice until processing.

### Tissue Processing

#### Blood

This protocol is best performed in batches of 6 to 8 mice, staggering if more than 6-8 mice is necessary.

1. Transfer collected blood to a 5 mL Eppendorf tube and add ACK lysis buffer at a 1:4 blood:lysis buffer ratio and incubate on an end-over-end turner for 10 minutes.
2. Centrifuge at 1500 xg for 10 minutes at 4 degrees C.
3. Aspirate off the supernatant.
4. Resuspend pellet in 1 mL complete RPMI. Store on ice until ready to stain.

#### Bone Marrow

This protocol is best performed in batches of 6 to 8 mice, staggering if more than 6-8 mice is necessary.

1. Resuspend marrow in 200 μL PBS.
2. Add 800 μL ACK lysis and incubate on an end-over-end turner for 10 minutes.
3. Centrifuge at 1500 xg for 10 minutes at 4 degrees C.
4. Aspirate off the supernatant.
5. Resuspend in 3 mL complete RPMI. Store on ice until ready to stain.

#### Lung

This protocol is best performed in batches of 6 to 8 mice, staggering if more than 6-8 mice is necessary.

1. Prepare 2.5 mL of room temperature tumor digestion medium (per mouse) by combining the following:
  a. 250 μL Collagenase/Hyaluronidase
  b. 375 μL DNase I solution at 1 mg/mL
  c. 1.875 mL complete RPMI 1640 media
2. Transfer stored tissue to a petri dish. Mince up tissue into small pieces with a straight edge razor blade using a chopping motion. Use a fresh dish and blade per sample.

1. Rinse each dish with 2.5 mL of digestion medium prepared in step 1, and transfer the tissue and digestion medium to a 14 mL round-bottom tube.
2. Incubate at 37 degrees C for 20 minutes with shaking.
3. While sample is incubating, prepare a 70-micron nylon mesh strainer over a 50 mL conical tube and prime the strainer with 5 mL of complete RPMI.
4. Following incubation, transfer the digested tissue into the strainer and push digested tissue through the strainer using the rubber end of a 5 mL syringe plunger. Rinse with 25 mL complete RPMI. Push as much as possible of the remainder of the tissue though the strainer and rinse with an additional 20 mL of complete RPMI. (50 mL total). Use a fresh strainer and plunger per sample.
5. Centrifuge at 300 xg for 10 minutes at room temperature, using a slow deceleration setting. Carefully pour off and discard the supernatant.
6. Resuspend in 3 mL of ACK lysis buffer. Let sit at room temperature for 3 minutes before topping off with 47 mL of complete RPMI. (50 mL total).
7. Centrifuge at 300 xg for 10 minutes at room temperature, using a slow deceleration setting. Carefully pour off and discard the supernatant.
8. Resuspend in 2 mL of complete RPMI. Store on ice until ready to stain.

### Flow Cytometry

This protocol is best performed in batches of 6 to 8 mice, staggering if more than 6-8 mice is necessary.

1. Stain appropriate samples with Vybrant DyeCycle Violet using the following amounts:
  a. Blood: 1 μL/mL
  b. Bone Marrow: 5 μL/mL
  c. Lung: 5 μL/mL
  d. Control cells: 1 μL/1 mL (using a maximum concentration of 1,000,000 cells/mL in complete RPMI)
2. Transfer to FACS tubes through a 40 micron cap filter and incubate for 30 min at 37 degrees C protected from light. Store on ice, protected from light until ready to flow.
3. Flow cells, with the following considerations:
  a. Controls:
    I. PC3-GFP-Luc cells, Unstained
    II. PC3-GFP-Luc cells, Stained with Vybrant Dye
    III. Cycle Violet
    IV. Matched tissue (blood, bone marrow, and/or lung) from an uninjected mouse, Unstained
    V. Matched tissue (blood, bone marrow, and/or lung) from an uninjected mouse, Stained with Vybrant DyeCycle Violet
    VI. Matched tissue (blood, bone marrow, and/or lung) from uninjected mouse, spiked with PC3-GFP-Luc cells, Unstained
    VII. Matched tissue (blood, bone marrow, and/or lung) from uninjected mouse, spiked with PC3-GFP-Luc cells, Stained with Vybrant DyeCycle
    VIII. When appropriate, PC3-GFP-Luc High-Ploidy, Unstained
    IX. When appropriate, PC3-GFP-Luc High-Ploidy, Stained with Vybrant DyeCycle Violet
    X. When appropriate, Matched tissue (blood, bone marrow, and/or lung) from uninjected mouse, spiked with PC3-GFP-Luc High-Ploidy cells, Unstained
    XI. When appropriate, Matched tissue (blood, bone marrow, and/or lung) from uninjected mouse, spiked with PC3-GFP-Luc High-Ploidy cells, Stained with Vybrant DyeCycle
  b. Run conditions:
    I. Run on Thermo-Fisher’s Attune NxT Acoustic Focusing Cytometer using the following modifications to accommodate analysis of a broader range of cell sizes:
      i. Hardware: Largest commercially available blocker bar installed over laser
      ii. Hardware: Alternative optical configuration. (Supplemental Figure 1).
      iii. Software: Use of SSC for thresholding, rather than standard FSC.
      iv. Software: 0.6 Area scaling factor for all lasers, rather than standard area scaling factors.
    II. Optimize voltages for SSC, VL1-A, VL2-A, and BL1-A channels.
    III. Run at least 50,000 events for control samples at 200 μL per minute. Run entire volume of blood, bone, and lung tissue samples at 1000 μL per minute.

### Data Analysis

1. Compensation Strategy
  a. BLI-A
    I. Positive: GFP+ Signal among “Matched tissue (blood, bone marrow, and/or lung) from uninjected mouse, spiked with PC3-GFP-Luc cells, Unstained” control
    II. Negative: “Matched tissue (blood, bone marrow, and/or lung) from uninjected mouse, Unstained” control
  b. VL2-A
    I. Positive: “Matched tissue (blood, bone marrow, and/or lung) from uninjected mouse, Stained with Vybrant DyeCycle Violet” control
    II. Negative: “Matched tissue (blood, bone marrow, and/or lung) from uninjected mouse, Unstained” control
2. Gating Strategy
  a. Gate 1a: SSC-A linear vs. VL1-A linear (Size-based exclusion)
  b. Gate 1b: SSC-A log vs. VL1-A log (Size-based exclusion)
  c. Gate 2: SSC-A linear vs. SSC-H linear (Doublet exclusion)
  d. Gate 3: Comp-VL2-A Histogram (Vybrant DyeCycle Violet unstained exclusion)
  e. Gate 4: Comp-BL1-A Histogram (GFP-negative exclusion)

## Results

We highlight the successful co-identification of CTCs and bone-marrow DTCs from a single tumor-bearing mouse using a flow-cytometry approach. The identification of rare events (here, CTCs and DTCs) by flow cytometry requires a stringent analysis approach. We developed and confirmed the following strict gating strategy.

Gate 1a is a size-exclusion gate defined by the distribution of control PC3-GFP-Luc cells stained with Vybrant DyeCycle Violet on an SSC-A linear vs. VL1-A linear density plot (Figure 2A). Gate 1a is then either confirmed or minutely adjusted into size-exclusion gate 1B via the distribution of control PC3-GFP-Luc cells stained with Vybrant DyeCycle Violet on a SSC-A log vs. VL1-A log density plot (Figure 2B). Together, these gates eliminate contaminating small debris and large, irregularly shaped foreign matter. Gate 2 is a doublet-exclusion gate defined by the distribution of control PC3-GFP-Luc cells stained with Vybrant DyeCycle Violet on an SSC-A linear vs. SSC-H linear density plot (Figure 2C). This gate eliminates doublet pairs of cells that are bound together. Gate 3 is a Vybrant DyeCycle Violet unstained exclusion gate defined by the distribution of control PC3-GFP-Luc cells stained with Vybrant DyeCycle Violet on a Comp-VL2-A histogram plot (Figure 2D). Gate 3 is applied to high-ploidy cells in Supplemental Figure 4D. This gate eliminates residual red blood cells that lack nuclear content and thus are not amenable to Vybrant DyeCycle Violet staining, as well as apoptotic or other dead or dying cells that fail to be stained by this live cell dye.

**Figure 2:**
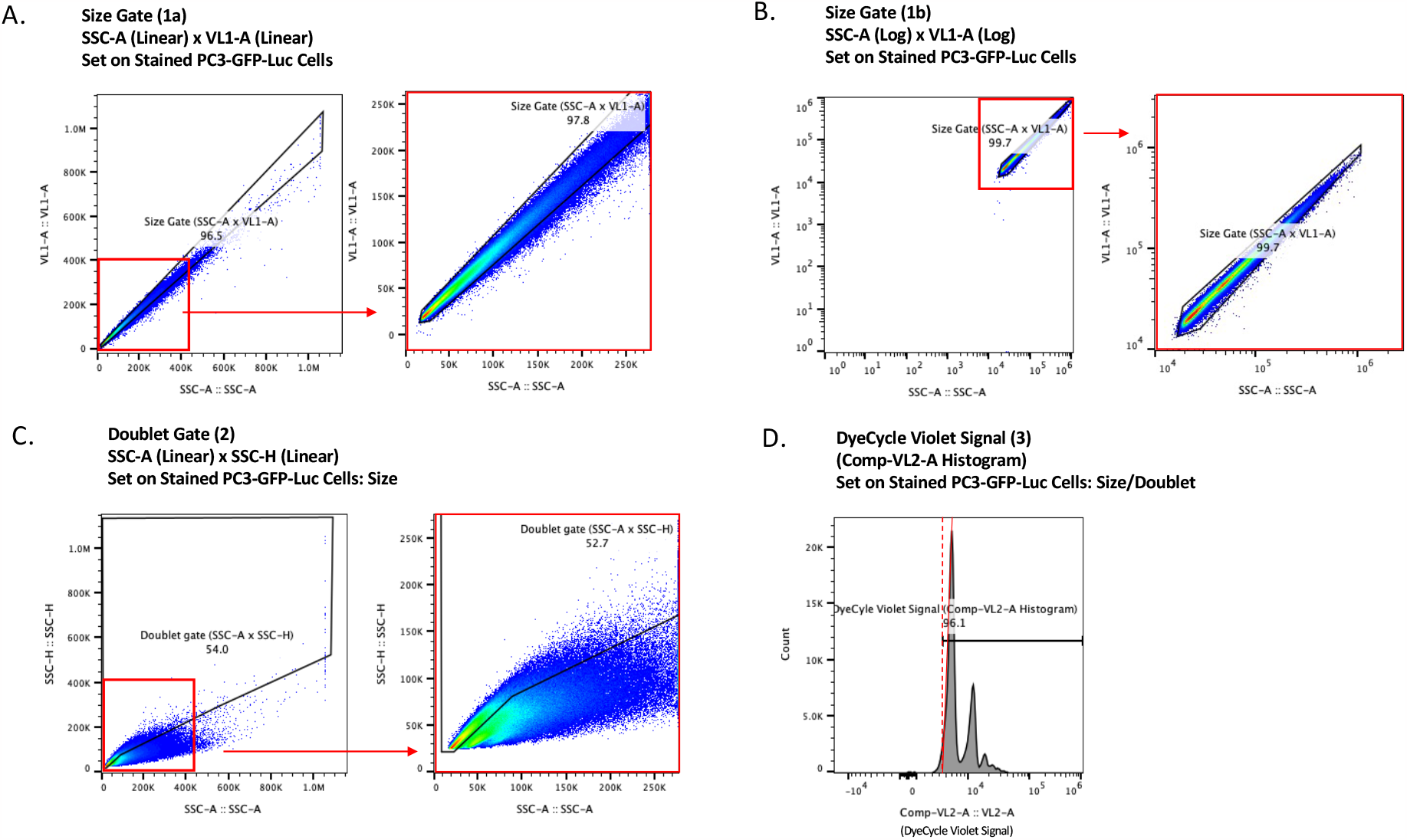
Pan-Tissue Gating (Gates 1-3) A) Size-exclusion Gate 1a defined using the distribution of control PC3-GFP-Luc cells stained with Vybrant DyeCycle Violet on an SSC-A linear vs. VL1-A linear density plot. B) Size-exclusion Gate 1b defined using the distribution of control PC3-GFP-Luc cells stained with Vybrant DyeCycle Violet on an SSC-A log vs. VL1-A log density plot. C) Doublet-exclusion Gate 2 defined by the distribution of control PC3-GFP-Luc cells stained with Vybrant DyeCycle Violet on an SSC-A linear vs. SSC-H linear density plot. D) Vybrant DyeCycle Violet unstained exclusion Gate 3 defined by the distribution of control PC3-GFP-Luc cells stained with Vybrant DyeCycle Violet on a Comp-VL2-A histogram plot.

Gates 1-3 are considered pan-tissue, as they are established using control cultured cells, and are appropriate to apply to flow analyte sourced from any tissue. Gate 4 is a tissue-specific GFP-negative exclusion gate defined by the distribution of events analyzed in the uninjected control sample of tissue of interest stained with Vybrant DyeCycle Violet on a Comp-BL1-A histogram plot (Figure 3). Each tissue has a unique level of background auto-fluorescence that emitted into the BL1-A channel. It is critical to set this gate individually per tissue type, to the lowest intensities that still exclude 100% of events. Gate 4 is defined in blood samples in Figure 3A and in bone marrow samples in figure 3B. This gate determines the tissue-specific GFP autofluorescence threshold for reliable detection of GFP-positivity-defined CTCs and DTCs.

**Figure 3:**
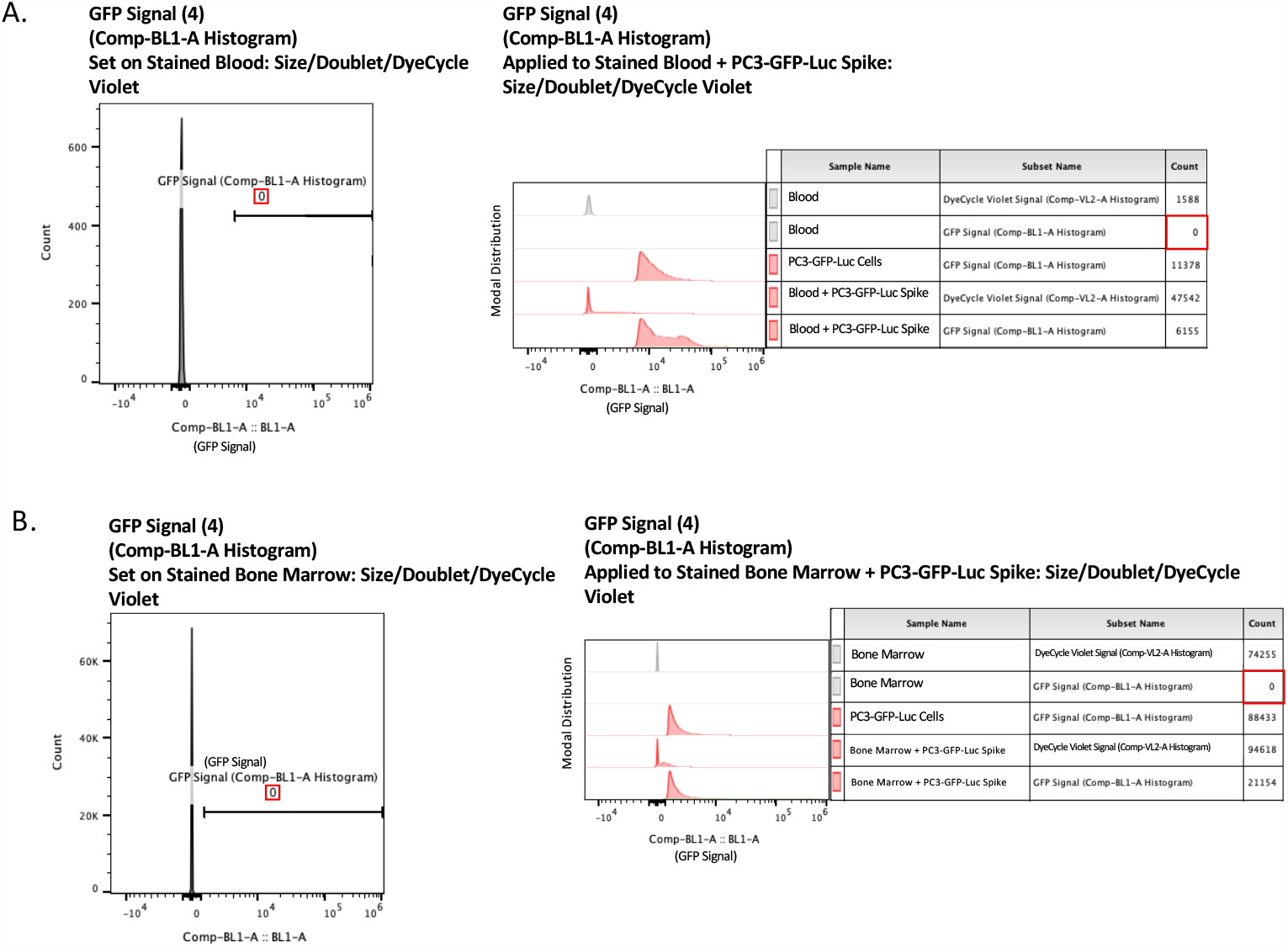
Tissue-Specific Gating (Gate 4) A) Tissue-specific GFP-negative exclusion Gate 4 defined by the distribution of events analyzed in the uninjected control sample of murine blood stained with Vybrant DyeCycle Violet on a Comp-BL1-A histogram plot, and then applied to uninjected murine blood samples that have been spiked with PC3-GFP-Luc cells and then stained with Vybrant DyeCycle Violet. B) Tissue-specific GFP-negative exclusion Gate 4 defined by the distribution of events analyzed in the uninjected control sample of murine bone marrow stained with Vybrant DyeCycle Violet on a Comp-BL1-A histogram plot, and then applied to uninjected murine bone marrow samples that have been spiked with PC3-GFP-Luc cells and then stained with Vybrant DyeCycle Violet.

The previously defined Pan Tissue Gates 1-3 are applied to blood and bone marrow samples that have been spiked with PC3-GFP-Luc cells and then stained with Vybrant DyeCycle Violet in Supplemental Figure 2. (Gate 1a applied in Supplemental Figure 2A, Gate 1b applied in Supplemental Figure 2B, Gate2 applied in Supplemental Figure 2C, and Gate 3 applied in Supplemental Figure 2D.) The previously defined tissue-specific Gate 4 is applied to blood and bone marrow samples that have been spiked with PC3-GFP-Luc cells and then stained with Vybrant DyeCycle Violet in Figure 3A and 3B, respectively.

Following the establishment of the gating approach, we optimized the upstream tissue processing conditions for ideal cellular recovery. To do this, we spiked 1,000,000 PC3-GFP-Luc normal or high-ploidy cells into blood or bone marrow harvested from mature adult male NSG mice and assayed various processing conditions including a) fixed cells vs. live cells, b) for fixed cells, permeabilized cells vs. unpermeabilized cells, and c) ACK lysed vs. Unlysed cells. For these conditions, fixed cell samples were fixed with 4% PFA at room temperature for 10 minutes and then washed 3 times with PBS, and permeabilized cell samples were permeabilized with 1% Saponin in PBS. We included high-ploidy cells in the optimization so as to ensure downstream applicability of this protocol to cells of various sizes. We found that live, ACK lysed cells (both normal and high-ploidy) could be best recovered from both blood and bone marrow (Figure 4).

**Figure 4:**
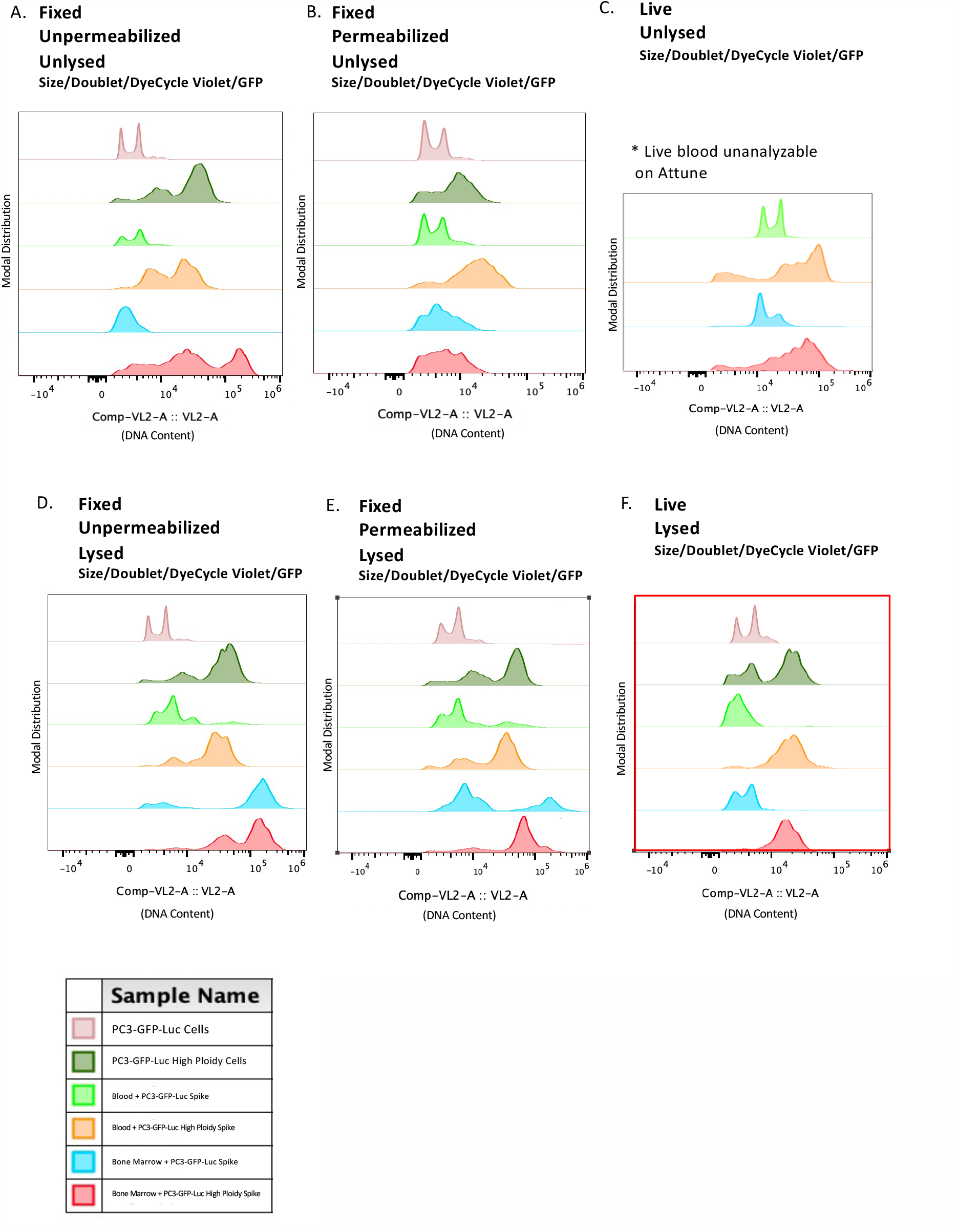
Analyte Preparation Optimization Comp-VL2-A histograms of PC3-GFP-Luc normal or high-ploidy cells alone, spiked into murine blood, or spiked into murine bone marrow, remaining following application of Gates 1-4. A) Cells were fixed, but not permeabilized nor lysed preceding cytometry. B) Cells were fixed and permeabilized, but not lysed preceding cytometry. C) Cells were not fixed, not permeabilized, and not lysed preceding cytometry. D) Cells were fixed, were not permeabilized, but were lysed preceding cytometry. E) Cells were fixed, permeabilized, and lysed preceding cytometry. F) Cells were not fixed nor permeabilized but were lysed preceding cytometry.

Following optimization of the tissue processing pipeline, we determined the sensitivity of the instrument, the Attune NxT Acoustic Focusing Flow Cytometer, to detect rare events. To do this, we spiked a limiting dilution (20,000; 10,000; 5,000; 1,000; 500; 250; 100; 50; 25; 10; or 0) of PC3-GFP-Luc cells into blood or bone marrow harvested from mature adult male NSG mice and measured the resultant detection (recovery) rate. In blood, even at low-input, spiked-in cells could be recovered when comprising as rare as 0.0019% of total sample analyzed (10 cells found within 525,546 total cells analyzed). In bone marrow, even at low input, spiked-in cells could be detected when comprising as rare as 0.0011% of sample (9 cells found within 829,431 cells analyzed) (Figure 5).

**Figure 5:**
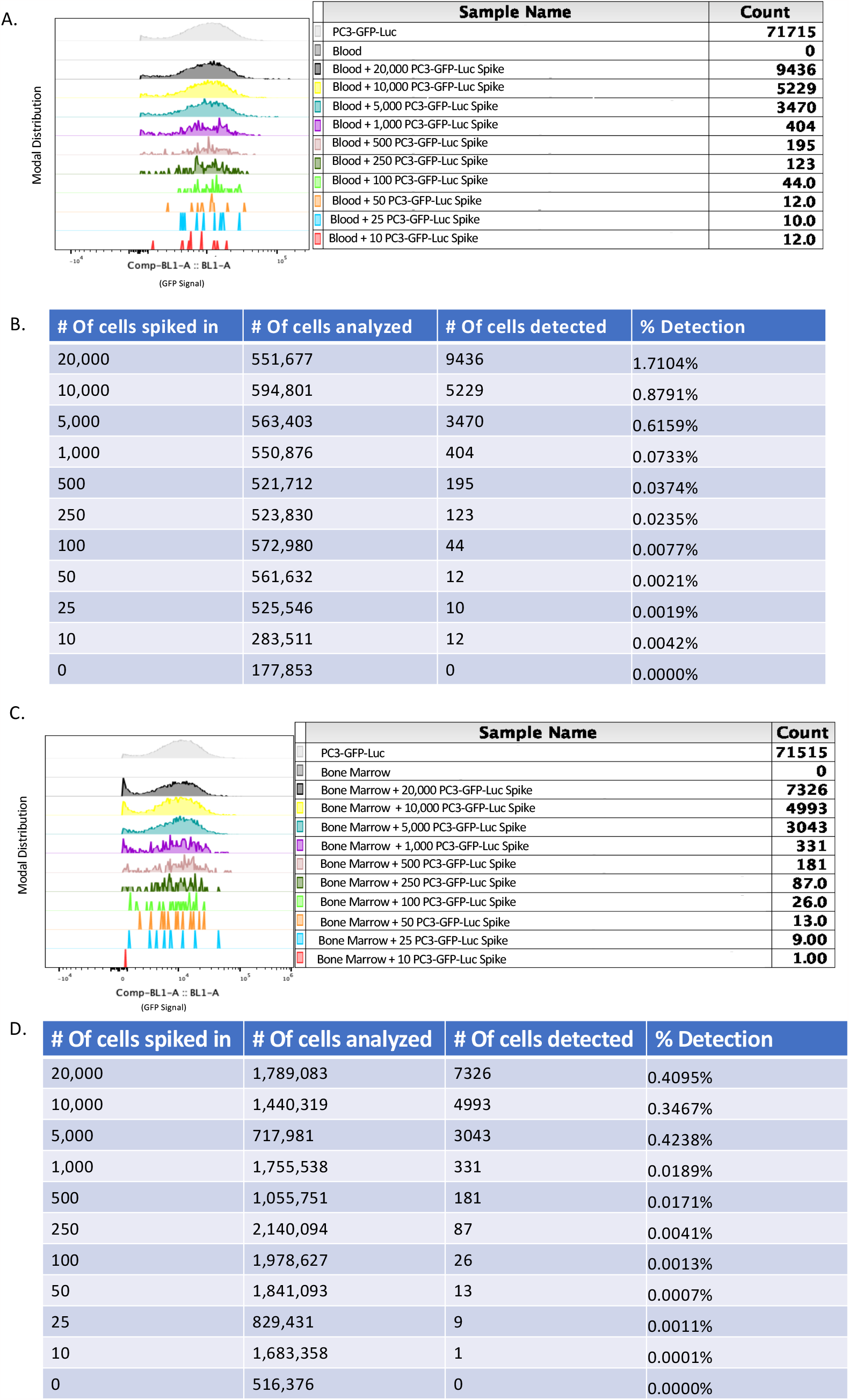
Limiting Dilution A) Comp-BL1-A Histogram of recovered PC3-GFP-Luc cells spiked into murine blood via limiting dilution. B) Percent detection of recovered PC3-GFP-Luc cells spiked into murine blood via limiting dilution. C) Comp-BL1-A Histogram of recovered PC3-GFP-Luc cells spiked into murine bone marrow via limiting dilution. D) Percent detection of recovered PC3-GFP-Luc cells spiked into murine bone marrow via limiting dilution.

In the example data showcased here, blood CTCs and bone DTCs are recovered using the aforementioned sample processing, data recording, and data analysis techniques 6 weeks after right-flank subcutaneous injection of PC3-GFP-Luc human prostate cancer cells into 10-week old male immunocompromised mice (Figure 6A). CTCs are identified in the blood of 2/8 injected mice (n=106 and 252 CTCs per mouse, respectively) and 0/4 mock-injected mice (Figure 6B). DTCs are identified in the bone marrow of 8/8 injected mice (n=1 DTC in 7 mice and 60 DTCs in 1 mouse) and 0/4 mock-injected mice (Figure 6 A).

**Figure 6:**
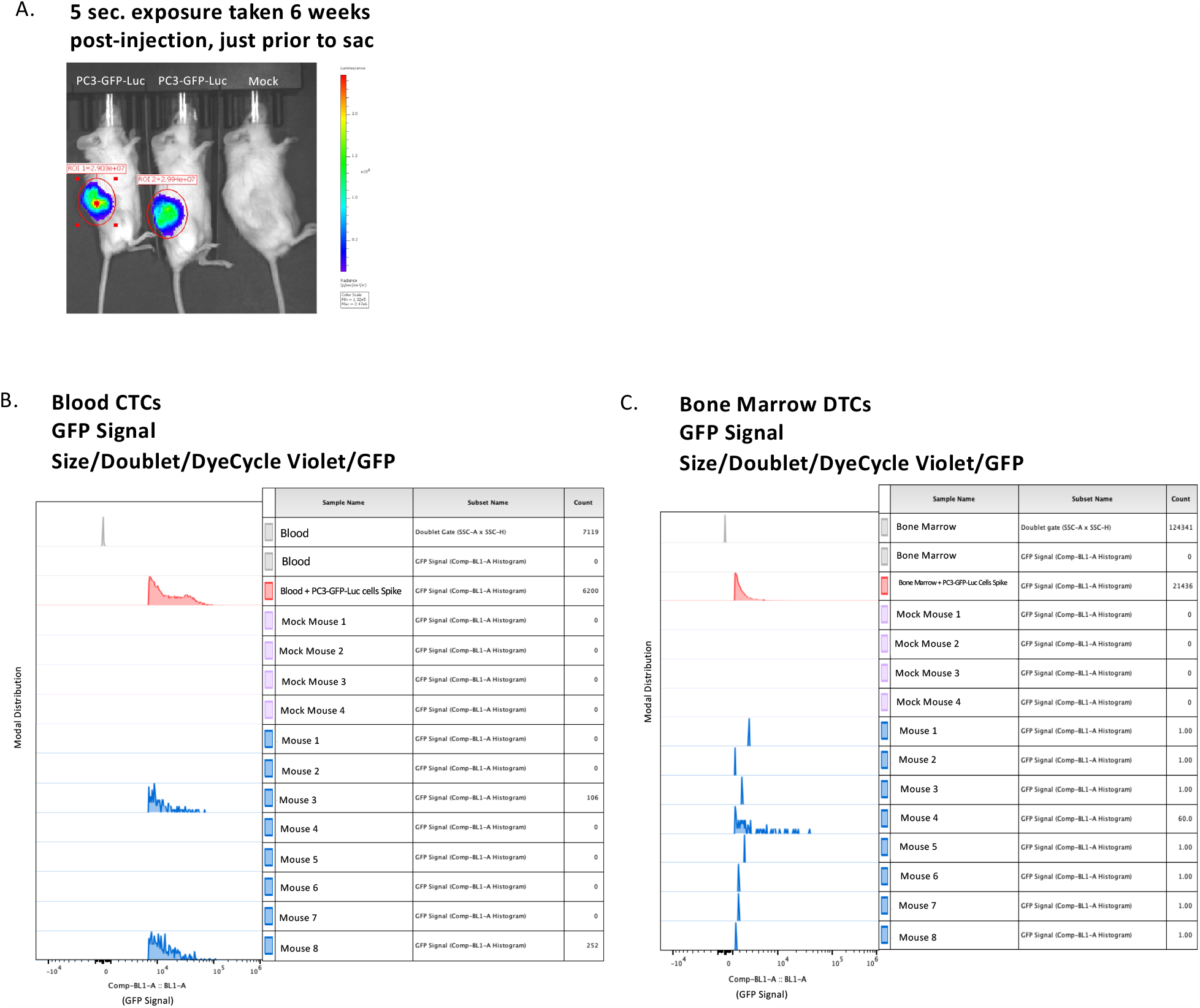
CTCs and DTCs Recovered from Blood and Bone Marrow Following Subcutaneous Injection A) Representative BLI image of mice 6 weeks after subcutaneous injection of PC3-GFP-Luc cells or mock. B) Comp-BL1-A histogram of blood-recovered CTCs from a subcutaneous metastasis model following application of Gates 1-4. C). Comp-BL1-A histogram of bone-marrow recovered DTCs from a subcutaneous metastasis model following application of Gates 1-4.

Additional data exemplifies the applicability of this protocol to a) cultured cells of alterative sizes and b) alternative harvested tissues. We include use of high-ploidy cells to demonstrate that this system is also appropriate for large cells. To make certain that Gate 1 is appropriate for larger high-ploidy cells, the axis should be adjusted so inclusion of the northwest corner of the density plot is ensured. Gates 1a and 1b are applied to high-ploidy cells alone and high-ploidy cells spiked into blood and bone marrow in Supplemental Figures 4A and 4B. To make certain that Gate 2 is appropriate for larger high-ploidy cells, the axis should be adjusted so inclusion of the western edge of the density plot is ensured. Gate 2 is applied to high-ploidy cells alone and high-ploidy cells spiked into blood and bone marrow in Supplemental Figure 4C. Various tissue-specific Gate 4s are applied to high-ploidy cells alone and high-ploidy cells spiked into blood and bone marrow in Supplemental Figure 4E. Furthermore, we include use of a tail-vein direct to lung injection model to demonstrate that this system is amenable to recovery of various organ types. Gates 1-3 are applied to lung samples that have been spiked with PC3-GFP-Luc cells and then stained with Vybrant DyeCycle Violet in Supplemental Figure 3A-D. Gate 4 is defined using an uninjected control sample of lung tissue stained with Vybrant DyeCycle Violet on a Comp-BL1-A histogram plot in Supplemental Figure 3E and then applied to lung samples that have been spiked with PC3-GFP-Luc cells and then stained with Vybrant DyeCycle Violet in Supplemental Figure 3F. Combined detection of a) high-ploidy cells in b) lung tissue is highlighted in Supplemental Figure 4A-E, in which previously defined Gates 1-4 are applied to high ploidy cells spiked into lung tissue.

In a second example, lung DTCs are recovered 3 days after tail vein injection of either normal or high-ploidy PC3-GFP-Luc cells into 10-week old male immunocompromised mice (Supplemental Figure 5A and 5B). DTCs were identified in 4/4 mice injected with normal cells and in 4/4 mice injected with high-ploidy cells. All mice had either 1 or 2 DTCs (Supplemental Figure 5C). Furthermore, an additional analysis of the ploidy of the identified DTCs shows that all DTCs recovered in mice injected with normal cancer cells displayed a normal amount of genomic content, while all DTCs recovered in mice injected with high-ploidy cancer cells displayed an increased amount of genomic content, consistent with their high ploidy status (Supplemental Figure 5D). This additional analysis was performed using the Vybrant DyeCycle Violet stain and typifies the kind of supplemental analyses that can be complexed with this approach for further CTC and DTC characterization.

## Discussion

Rapid and reliable CTC and DTC detection forms a major underpinning of rigorous *in vivo* metastasis research. While many cancer cells initiate metastasis, very few can complete it. Many motile or invasion metastasis-initiating cells never thrive as CTCs, and many CTCs never thrive as DTCs. Each successive step of metastatic cascade presents increasing barriers to metastatic success. As such, there is need for metastasis research approaches that provide a more detailed understanding of the specific failure-points of various metastasis models along the metastatic cascade. Here, we present a novel method that allows for the simultaneous comparison of multiple steps of the cascade within one model system via the identification and phenotyping of CTCs and DTCs from a single animal.

This experimental design is preferred because it eliminates inappropriate cross-cascade inference-making. For example, observation of increased numbers of CTCs in the circulation cannot be used to independently predict future increased metastatic burden. Without orthogonal DTC data, the CTC data alone remains too limited to assess the presence (or absence) of a possible extravasation-incompetency bottleneck.

Using this approach, we detected abundant CTCs within 500 mL of blood from 2/8 animals. We assume the lack of detected CTCs in the other 6 mice was due to biological differences such as slower subcutaneous tumor growth rate, rather than instrument failure (i.e., it is likely that those 6 animals did not have CTCs at time of monitoring). Additionally, we were able to detect both bone and lung DTCs in every animal analyzed. The limited number of DTCs identified in comparison to the number of CTCs aligns with clinical reports of an extravasation-bottleneck. In one animal, 60 bone marrow DTCs were identified, indicating likely early colonization of a DTC into a micro-metastatic lesion as yet too small to be visualized via bioluminescent imaging prior to the experimental end-point.

This specific protocol was developed and optimized for use with Thermo-Fisher’s Attune NxT Acoustic Focusing Cytometer equipped with the following four lasers: Violet (405nm), Blue (488nm), Yellow (561 nm), and Red (637 nm). Using this machine, our method is incredibly amenable to adaptation. It is compatible with wide range of cell sizes and can be complexed with other fluorescent-based markers to further characterize recovered CTCs and DTCs. For example, we have successfully analyzed physically enlarged high-ploidy cells, adding a violet fluorescent DNA content dye to obtain a confirmatory read-out of ploidy. Any other markers of interest could easily be added, assuming appropriate optical channel diversity. Additionally, it is compatible with other injection techniques and analysis of other secondary organs including solid organs, such as lung tissue recovery following tail vein injection.

Existing CTC and DTC detection methodologies are comparatively laborious than this flow cytometry approach. We have previously optimized and published a robust CTC and DTC detection immunofluorescence-imaging based method ((doi: 10.1186/s12575-018-0078-5; doi: 10.18632/oncotarget.12000). This imaging-based technique requires smearing of harvested cells on slides, followed by mouse vs. human cell-specific differential immunofluorescent staining and time-intensive slide scanning using the Metafer classifier detection system. The abundance of cells in the bone marrow collected from two hind limbs (femur and tibia) of an average adult male mouse frequently required more than 8 slides, the single run-maximum limit of the Metafer detection system. Considering an 8-slide scan time of 12 hours, it would take upwards of one month to scan a single experiment’s worth of slides. Furthermore, though the automated classifier-based detection system was effective in identifying potential suspected human-origin CTC or DTCs among murine cells based on differential fluorescence, human-verification was still required to remove about 30% of called events as false-positives.

Our new flow-cytometry-based technique reduces the data collection time from a month or more to several hours, creating possibility for higher-powered experiments with increased numbers of animals. Additionally, it allows for the simultaneous processing of additional murine tissues per animal, such as dissociated lungs, thus increasing both the breadth and depth of collectable data.

Some potential limitations of our system include the reliance on strong, constitutive, intracellular fluorescent protein expression. Though non-concerning in immunocompromised animal models, the immunoreactivity of GFP has been reported in immunocompetent model systems. Also, while most of the flow cytometer hardware and software settings that we used to achieve these results are temporary and easily exchangeable, the required installation of the largest commercially available blocker bar over our machine’s internal laser system is a permanent modification. We have not tested the reliability of this approach using a machine that does not contain the blocker bar, though we hypothesize it would severely limit the ability to detect larger sized CTC and DTC events. A potential improvement to this protocol would be apply this approach to imaging flow cytometry to simultaneously obtain imaging data of detected CTCs and DTCs that could further inform experiments.

