## Supplemental for "Multiparameter flow cytometric detection and analysis of rare cells *in in vivo* models of cancer metastasis"

### Supplemental Figure 1: Optical Configurations

A. **Attune NxT Standard Optical Configuration (Performance Test):** B. **Attune NxT Adjusted Optical Configuration (Experimental Samples):**

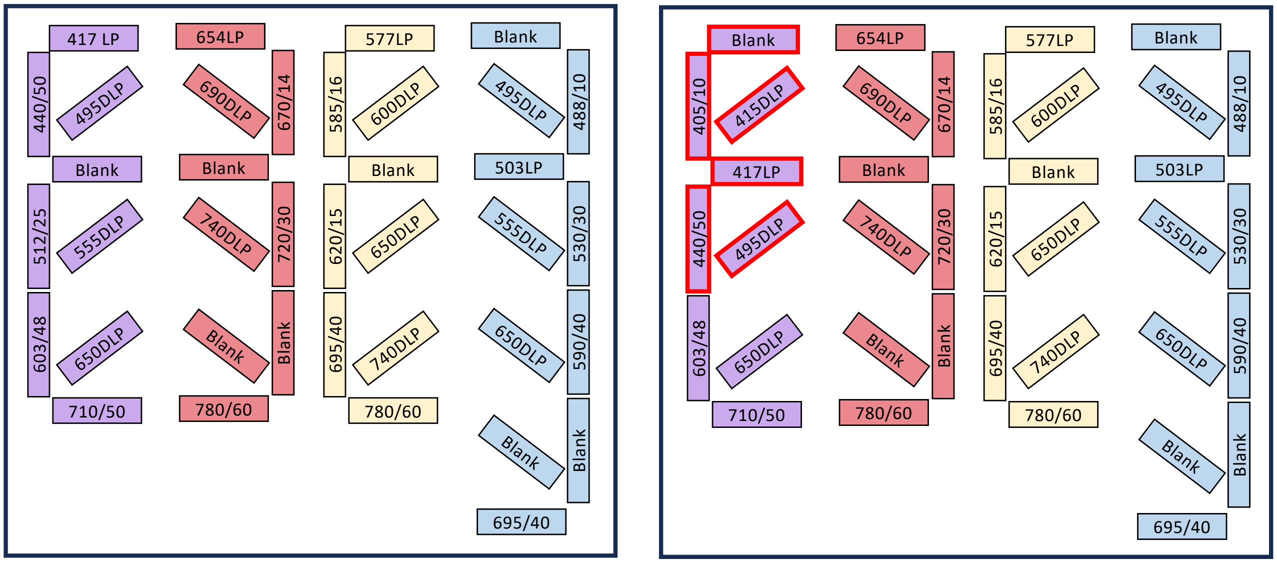


Supplemental Figure 1: A) Standard optical configuration for Attune NxT Acoustic Focusing Cytometer used during the "performance test" step of daily instrument set-up. B) Modified optical configuration for Attune NxT Acousting Focusing Cytometer used for outlined CTC and DTC detection protocol.

### Supplemental Figure 2: Pan-Tissue Gating (Gates 1-3) Applied to Spiked Blood and Bone Marrow Samples

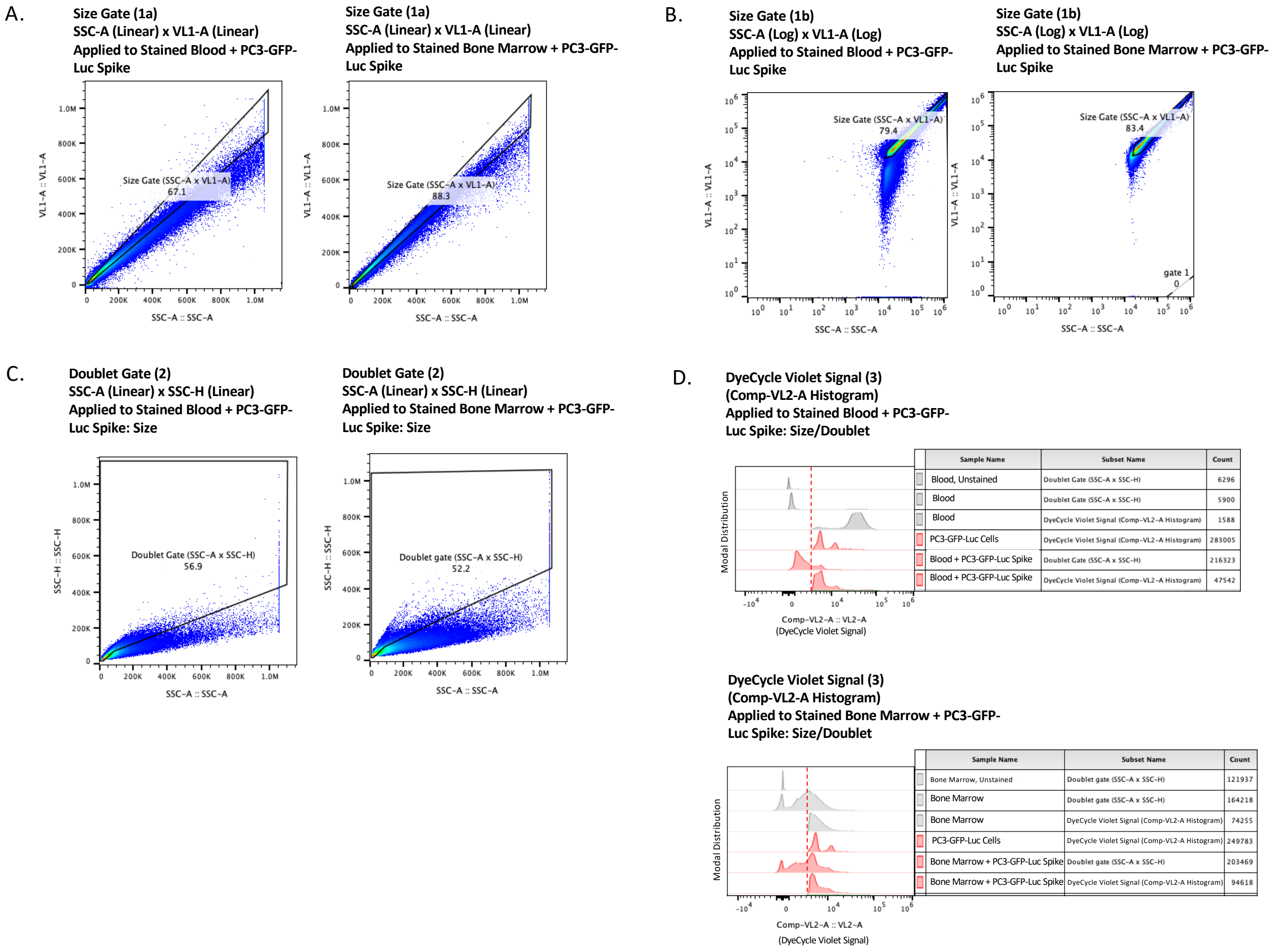


| Sample Name | Subset Name | Count |
|---|---|---|
| Blood, Unstained | Doublet Gate (SSC-A x SSC-H) | 6296 |
| Blood | Doublet Gate (SSC-A x SSC-H) | 5900 |
| Blood | DyeCycle Violet Signal (Comp-VL2-A Histogram) | 1588 |
| PC3-GFP-Luc Cells | DyeCycle Violet Signal (Comp-VL2-A Histogram) | 283005 |
| Blood + PC3-GFP-Luc Spike | Doublet Gate (SSC-A x SSC-H) | 216323 |
| Blood + PC3-GFP-Luc Spike | DyeCycle Violet Signal (Comp-VL2-A Histogram) | 47542 |

| Sample Name | Subset Name | Count |
|---|---|---|
| Bone Marrow, Unstained | Doublet gate (SSC-A x SSC-H) | 121937 |
| Bone Marrow | Doublet gate (SSC-A x SSC-H) | 164218 |
| Bone Marrow | DyeCycle Violet Signal (Comp-VL2-A Histogram) | 74255 |
| PC3-GFP-Luc Cells | DyeCycle Violet Signal (Comp-VL2-A Histogram) | 249783 |
| Bone Marrow + PC3-GFP-Luc Spike | Doublet gate (SSC-A x SSC-H) | 203469 |
| Bone Marrow + PC3-GFP-Luc Spike | DyeCycle Violet Signal (Comp-VL2-A Histogram) | 94618 |

Supplemental Figure 2: A) Size-exclusion Gate 1a applied to uninjected murine blood and bone marrow samples that have been spiked with PC3-GFP-Luc cells and then stained with Vybrant DyeCycle Violet. B) Size-exclusion Gate 1b applied to uninjected murine blood and bone marrow samples that have been spiked with PC3-GFP-Luc cells and then stained with Vybrant DyeCycle Violet. C) Doublet-exclusion Gate 2 applied to uninjected murine blood and bone marrow samples that have been spiked with PC3-GFP-Luc cells and then stained with Vybrant DyeCycle Violet. D) Vybrant DyeCycle Violet unstained exclusion Gate 3 histogram applied to uninjected murine blood and bone marrow samples that have been spiked with PC3-GFP-Luc cells and then stained with Vybrant DyeCycle Violet.

### Supplemental Figure 3: All Gating (Gates 1-4) Applied to Spiked Lung Samples

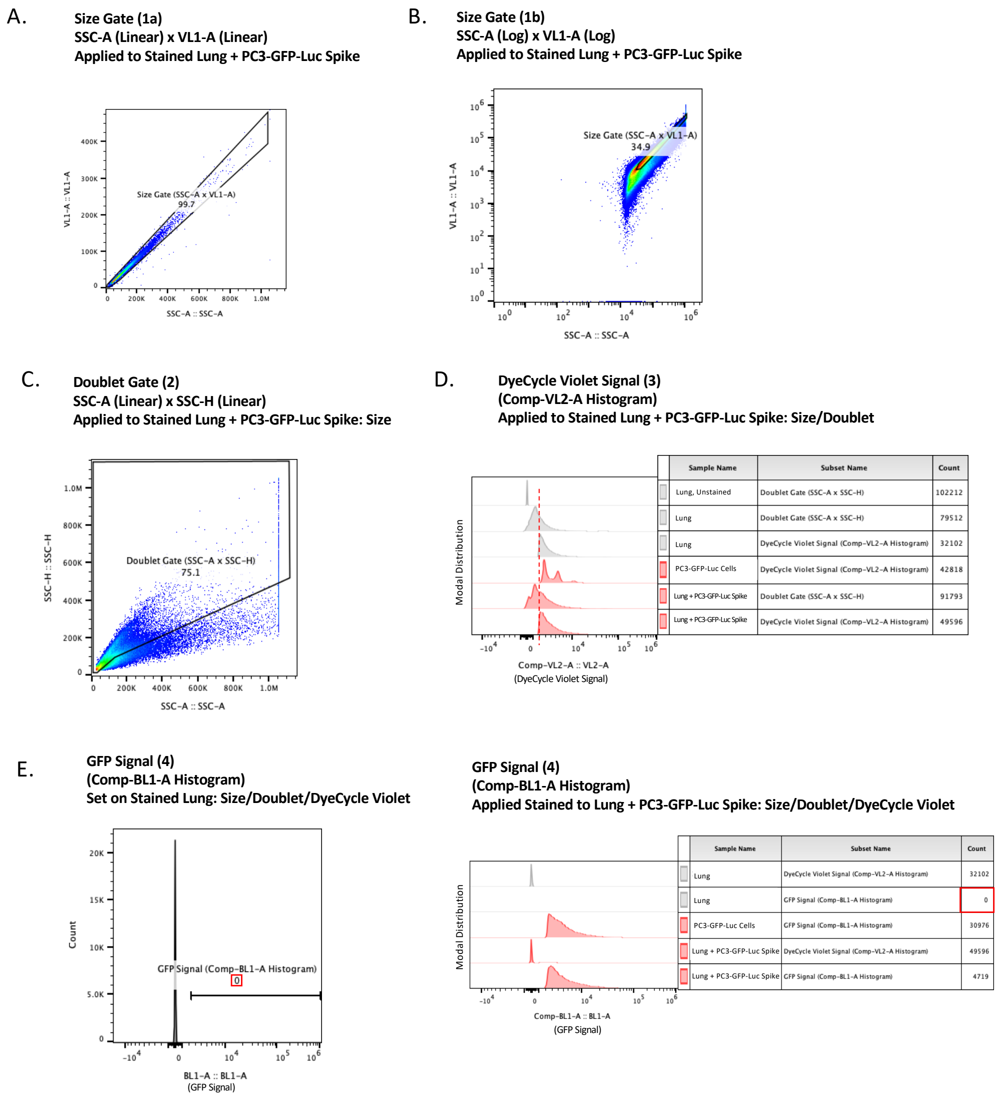


Supplemental Figure 3: A) Size-exclusion Gate 1a applied to uninjected murine lung tissue samples that have been spiked with PC3-GFP-Luc cells and then stained with Vybrant DyeCycle Violet. B) Size-exclusion Gate 1b applied to uninjected murine lung tissue samples that have been spiked with PC3-GFP-Luc cells and then stained with Vybrant DyeCycle Violet. C) Doublet-exclusion Gate 2 applied to uninjected murine lung tissue samples that have been spiked with PC3-GFP-Luc cells and then stained with Vybrant DyeCycle Violet. D) Vybrant DyeCycle Violet unstained exclusion Gate 3 histogram applied to uninjected murine lung tissue samples that have been spiked with PC3-GFP-Luc cells and then stained with Vybrant DyeCycle Violet. E) Tissue-specific GFP-negative exclusion Gate 4 defined by the distribution of events analyzed in the uninjected control sample of murine lung tissue stained with Vybrant DyeCycle Violet on a Comp-BL1-A histogram plot, and then applied to uninjected lung tissue samples that have been spiked with PC3-GFP-Luc cells and then stained with Vybrant DyeCycle Violet.

### Supplemental Figure 4: All Gating (Gates 1-4) Applied to High Ploidy Cell Types in Spiked Blood, Bone Marrow, and Lung Samples

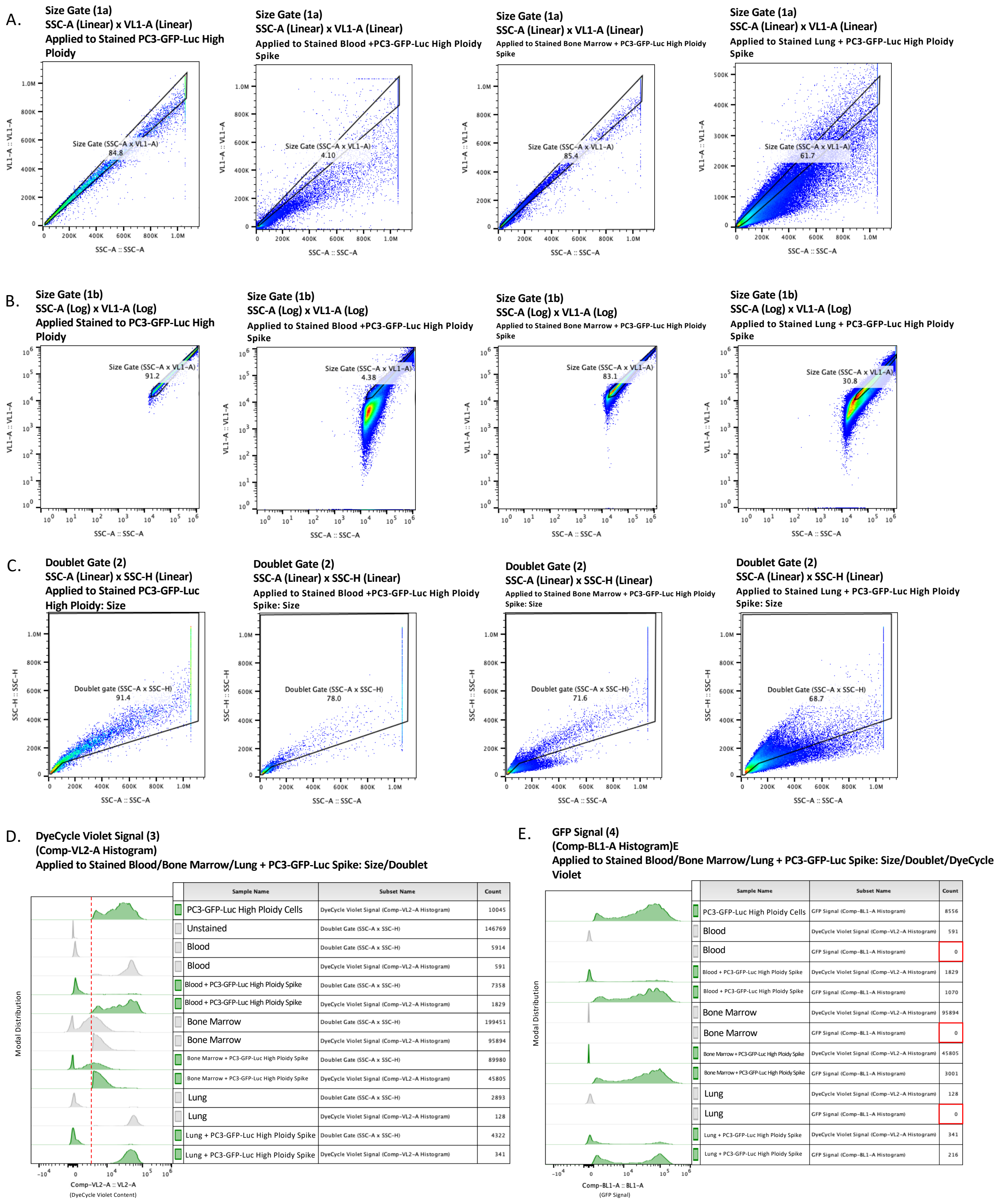


| Sample Name | Subset Name | Count |
|---|---|---|
| PC3-GFP-Luc High Ploidy Cells | DyeCycle Violet Signal (Comp-VL2-A Histogram) | 10045 |
| Unstained | Doublet Gate (SSC-A x SSC-H) | 146769 |
| Blood | Doublet Gate (SSC-A x SSC-H) | 5914 |
| Blood | DyeCycle Violet Signal (Comp-VL2-A Histogram) | 591 |
| Blood + PC3-GFP-Luc High Ploidy Spike | Doublet Gate (SSC-A x SSC-H) | 7358 |
| Blood + PC3-GFP-Luc High Ploidy Spike | DyeCycle Violet Signal (Comp-VL2-A Histogram) | 1829 |
| Bone Marrow | Doublet Gate (SSC-A x SSC-H) | 199451 |
| Bone Marrow | DyeCycle Violet Signal (Comp-VL2-A Histogram) | 95894 |
| Bone Marrow + PC3-GFP-Luc High Ploidy Spike | Doublet Gate (SSC-A x SSC-H) | 89980 |
| Bone Marrow + PC3-GFP-Luc High Ploidy Spike | DyeCycle Violet Signal (Comp-VL2-A Histogram) | 45805 |
| Lung | Doublet Gate (SSC-A x SSC-H) | 2893 |
| Lung | DyeCycle Violet Signal (Comp-VL2-A Histogram) | 128 |
| Lung + PC3-GFP-Luc High Ploidy Spike | Doublet Gate (SSC-A x SSC-H) | 4322 |
| Lung + PC3-GFP-Luc High Ploidy Spike | DyeCycle Violet Signal (Comp-VL2-A Histogram) | 341 |

| Sample Name | Subset Name | Count |
|---|---|---|
| PC3-GFP-Luc High Ploidy Cells | GFP Signal (Comp-BL1-A Histogram) | 8556 |
| Blood | DyeCycle Violet Signal (Comp-VL2-A Histogram) | 591 |
| Blood | GFP Signal (Comp-BL1-A Histogram) | 0 |
| Blood + PC3-GFP-Luc High Ploidy Spike | DyeCycle Violet Signal (Comp-VL2-A Histogram) | 1829 |
| Blood + PC3-GFP-Luc High Ploidy Spike | GFP Signal (Comp-BL1-A Histogram) | 1070 |
| Bone Marrow | DyeCycle Violet Signal (Comp-VL2-A Histogram) | 95894 |
| Bone Marrow | GFP Signal (Comp-BL1-A Histogram) | 0 |
| Bone Marrow + PC3-GFP-Luc High Ploidy Spike | DyeCycle Violet Signal (Comp-VL2-A Histogram) | 45805 |
| Bone Marrow + PC3-GFP-Luc High Ploidy Spike | GFP Signal (Comp-BL1-A Histogram) | 3001 |
| Lung | DyeCycle Violet Signal (Comp-VL2-A Histogram) | 128 |
| Lung | GFP Signal (Comp-BL1-A Histogram) | 0 |
| Lung + PC3-GFP-Luc High Ploidy Spike | DyeCycle Violet Signal (Comp-VL2-A Histogram) | 341 |
| Lung + PC3-GFP-Luc High Ploidy Spike | GFP Signal (Comp-BL1-A Histogram) | 216 |

Supplemental Figure 4: A) Size-exclusion Gate 1a applied to uninjected murine blood, bone marrow, and lung samples that have been spiked with PC3-GFP-Luc high-ploidy cells and then stained with Vybrant DyeCycle Violet. B) Size-exclusion Gate 1b applied to uninjected murine blood, bone marrow, and lung samples that have been spiked with PC3-GFP-Luc high-ploidy cells and then stained with Vybrant DyeCycle Violet. C) Doublet-exclusion Gate 2 applied to uninjected murine blood, bone marrow, and lung samples that have been spiked with PC3-GFP-Luc high-ploidy cells and then stained with Vybrant DyeCycle Violet. D) Vybrant DyeCycle Violet unstained exclusion Gate 3 applied to uninjected

### Supplemental Figure 5: DNA Content Analysis of High-Ploidy DTCs Recovered from Lung Tissue Following Tail-Vein Injection

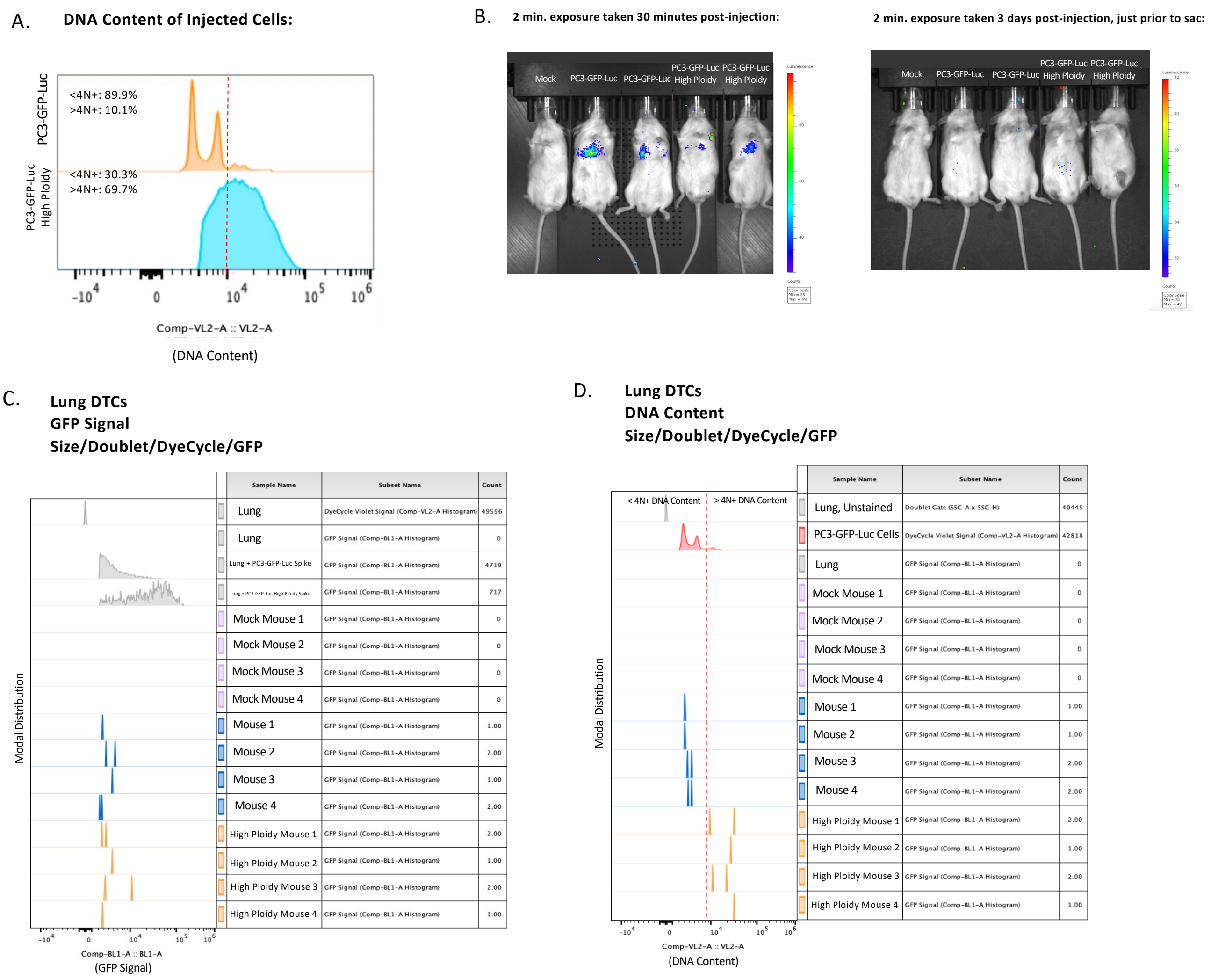


Supplemental Figure 5:
A) Ploidy status of injected PC3-GFP-Luc normal and high-ploidy cells at time of injection. B) Representative BLI image of mice within 30 minutes of tail-vein injection of PC3-GFP-Luc normal or high-ploidy cells, and again 3 days following injection. C) Comp-BL1-A histogram of lung tissue recovered DTCs from a tail vein metastasis model following application of Gates 1-4. D) Comp-VL2-A histogram of the lung tissue recovered DTCs used to measure DTC DNA content.
